## Supplementary material for "Effectiveness of *Pseudomonas aeruginosa* type VI secretion system relies on toxin potency and type IV pili-dependent interaction": S1 Table

**S1 Table List of strains used in this study**

| **Strain** | **Features** | **Source** |
| --- | --- | --- |
| PAO1 | Wild type strain, parental strain used in constructuin of all other strains in this study | Laboratory collection |
| ΔrsmN | Deletion of rsmN (PA5183.1) | This study |
| ΔrsmA | Deletion of rsmA (PA0905) | Laboratory collection |
| ΔretS | Deletion of retS (PA4856) | Laboratory collection |
| ΔrsmA ΔrsmN | Deletion of rsmA (PA0905) and rsmN (PA5183.1) | This study |
| ΔtssB1 ΔtssB2 ΔtssB3 | Deletion of rsmN (PA5183.1), tssB1 (PA0083), tssB2 (PA1657) and tssB3 (PA2365) | Laboratory collection |
| ΔrsmN ΔtssB1 ΔtssB2 ΔtssB3 | Deletion of tssB1 (PA0083), tssB2 (PA1657) and tssB3 (PA2365) | This study |
| ΔrsmA ΔtssB1 ΔtssB2 ΔtssB3 | Deletion of rsmA (PA0905), tssB1 (PA0083), tssB2 (PA1657) and tssB3 (PA2365) | Laboratory collection |
| ΔretS ΔtssB1 ΔtssB2 ΔtssB3 | Deletion of retS (PA4856), tssB1 (PA0083), tssB2 (PA1657) and tssB3 (PA2365) | This study |
| ΔrsmA ΔrsmN ΔtssB1ΔtssB2ΔtssB3 | Deletion of rsmA (PA0905), rsmN (PA5183.1), tssB1 (PA0083), tssB2 (PA1657) and tssB3 (PA2365) | This study |
| ΔrsmA tssB1:mScarlet-I | Deletion of rsmA (PA0905), C-terminal mScarlet-I fusion to tssB1 (PA0083) at native locus | Laboratory collection |
| ΔretS tssB1:mScarlet-I | Deletion of retS (PA4856), C-terminal mScarlet-I fusion to tssB1 (PA0083) at native locus | Laboratory collection |
| ΔrsmA ΔrsmN tssB1:mScarlet-I | Deletion of rsmA (PA0905) and rsmN (PA5183.1), C-terminal mScarlet-I fusion to tssB1 (PA0083) at native locus | This study |
| Δtse1_tsi1 | Deletion of tse1 and tsi1 (PA1844-45) | This study |
| Δtse2_tsi2 | Deletion of tse2 and tsi2 (PA2702-03) | This study |
| Δtse3_tsi3 | Deletion of tse3 and tsi3 (PA3484-85) | This study |
| Δtse4_tsi4 | Deletion of tse4 and tsi4 (PA2774-75) | This study |
| Δtse5_tsi5 | Deletion of tse5 and tsi5 (PA2683-84) | This study |
| Δtse6_tsi6 | Deletion of tse6 and tsi6 (PA1844-45) | This study |
| Δtse7_tsi7 | Deletion of tse7 and tsi7 (PA0099-0100) | This study |
| Δtse8_tsi8 | Deletion of tse8 and tsi8 (PA1844-45) | This study |
| ΔrsmA Δtse1_tsi1 | Deletion of rsmA (PA0905), tse1 and tsi1 (PA1844-45) | Laboratory collection |
| ΔrsmA Δtse2_tsi2 | Deletion of rsmA (PA0905), tse2 and tsi2 (PA2702-03) | Laboratory collection |
| ΔrsmA Δtse3_tsi3 | Deletion of rsmA (PA0905), tse3 and tsi3 (PA3484-85) | Laboratory collection |
| ΔrsmA Δtse4_tsi4 | Deletion of rsmA (PA0905), tse4 and tsi4 (PA2774-75) | This study |
| ΔrsmA Δtse5_tsi5 | Deletion of rsmA (PA0905), tse5 and tsi5 (PA2683-84) | This study |
| ΔrsmA Δtse6_tsi6 | Deletion of rsmA (PA0905), tse6 and tsi6 (PA1844-45) | This study |
| ΔrsmA Δtse7_tsi7 | Deletion of rsmA (PA0905), tse7 and tsi7 (PA0099-0100) | This study |
| ΔrsmA Δtse8_tsi8 | Deletion of rsmA (PA0905), tse8 and tsi8 (PA1844-45) | This study |
| ΔrsmA Δtse1_tsi1 Δtse3_tsi3 | Deletion of rsmA (PA0905), tse1 and tsi1 (PA1844-45), tse3 and tsi3 (PA3484-85) | Laboratory collection |
| ΔretS Δtse1_tsi1 | Deletion of retS (PA4856), tse1 and tsi1 (PA1844-45) | This study |
| ΔretS Δtse2_tsi2 | Deletion of retS (PA4856), tse2 and tsi2 (PA2702-03) | This study |
| ΔretS Δtse3_tsi3 | Deletion of retS (PA4856), tse3 and tsi3 (PA3484-85) | This study |
| ΔretS Δtse4_tsi4 | Deletion of retS (PA4856), tse4 and tsi4 (PA2774-75) | This study |
| ΔretS Δtse5_tsi5 | Deletion of retS (PA4856), tse5 and tsi5 (PA2683-84) | This study |
| ΔretS Δtse6_tsi6 | Deletion of retS (PA4856), tse6 and tsi6 (PA1844-45) | This study |
| ΔretS Δtse7_tsi7 | Deletion of retS (PA4856), tse7 and tsi7 (PA0099-0100) | This study |
| ΔretS Δtse8_tsi8 | Deletion of retS (PA4856), tse8 and tsi8 (PA1844-45) | This study |
| ΔrsmA ΔrsmN Δtse1_tsi1 | Deletion of rsmA (PA0905), rsmN (PA5183.1), tse1 and tsi1 (PA1844-45) | This study |
| ΔrsmA ΔrsmN Δtse2_tsi2 | Deletion of rsmA (PA0905), rsmN (PA5183.1), tse2 and tsi2 (PA2702-03) | This study |
| ΔrsmA ΔrsmN Δtse3_tsi3 | Deletion of rsmA (PA0905), rsmN (PA5183.1), tse3 and tsi3 (PA3484-85) | This study |
| ΔrsmA ΔrsmN Δtse4_tsi4 | Deletion of rsmA (PA0905), rsmN (PA5183.1), tse4 and tsi4 (PA2774-75) | This study |
| ΔrsmA ΔrsmN Δtse5_tsi5 | Deletion of rsmA (PA0905), rsmN (PA5183.1), tse5 and tsi5 (PA2683-84) | This study |
| ΔrsmA ΔrsmN Δtse6_tsi6 | Deletion of rsmA (PA0905), rsmN (PA5183.1), tse6 and tsi6 (PA1844-45) | This study |
| ΔrsmA ΔrsmN Δtse7_tsi7 | Deletion of rsmA (PA0905), rsmN (PA5183.1), tse7 and tsi7 (PA0099-0100) | This study |
| ΔrsmA ΔrsmN Δtse8_tsi8 | Deletion of rsmA (PA0905), rsmN (PA5183.1), tse8 and tsi8 (PA1844-45) | This study |
| ΔrsmA Δtle1_tli1ab | Deletion of rsmA (PA0905), tle1, tli1a and tli1b (PA3290-92) | This study |
| ΔrsmA Δtle3_tli3 | Deletion of rsmA (PA0905), tle3 and tli3 (PA0259-60) | This study |
| ΔrsmA Δtle4_tli4 | Deletion of rsmA (PA0905), tle4 and tli4 (PA1509-10) | This study |
| ΔrsmA ΔpldA_tli5a | Deletion of rsmA (PA0905), pldA and tli5a (PA3487-88) | This study |
| ΔrsmA ΔpldB_tli5b1-3 | Deletion of rsmA (PA0905), pldB, tli5b1, tli5b2 and tli5b3 (PA5086-89) | This study |
| ΔrsmA ΔtseT_tsiT | Deletion of rsmA (PA0905), tseT and tsiT (PA3907-08) | This study |
| ΔrsmA ΔtseV_tsiV | Deletion of rsmA (PA0905), tseV and tsiV (PA0821-22) | This study |
| ΔrsmA ΔvgrG2b_vgrG2bi | Deletion of rsmA (PA0905), vgrG2b and vgrG2bi (PA0261-62) | This study |
| ΔrsmA ΔampDh3_ampDh3i | Deletion of rsmA (PA0905), ampDh3 and ampDh3i (PA0807-08) | This study |
| ΔrsmA ΔPA5264_PA5265 | Deletion of rsmA (PA0905), PA5264 and PA5265 | This study |
| ΔrsmA Δazu | Deletion of rsmA (PA0905) and azu (PA4922) | This study |
| ΔretS Δtle1_tli1ab | Deletion of retS (PA4856), tle1, tli1a and tli1b (PA3290-92) | This study |
| ΔretS ΔpldA_tli5a | Deletion of retS (PA4856), pldA and tli5a (PA3487-88) | This study |
| ΔretS ΔpldB_tli5b1-3 | Deletion of retS (PA4856), pldB, tli5a, tli5b and tli5c (PA5086-89) | This study |
| ΔretS ΔtseT_tsiT | Deletion of retS (PA4856), tseT and tsiT (PA3907-08) | This study |
| ΔretS ΔampDh3_ampDh3i | Deletion of retS (PA4856), ampDh3 and ampDh3i (PA0807-08) | This study |
| ΔpilA | Deletion of pilA (PA4525) | This study |
| ΔrsmA ΔpilA | Deletion of rsmA (PA0905) and pilA (PA4525) | This study |
| ΔretS ΔpilA | Deletion of retS (PA4856) and pilA (PA4525) | This study |
| ΔrsmA ΔpilA ΔtssB1 ΔtssB2 ΔtssB3 | Deletion of rsmA (PA0905), pilA (PA4525), tssB1 (PA0083), tssB2 (PA1657) and tssB3 (PA2365) | This study |
| ΔrsmA ΔpilA ΔtseT_tsiT | Deletion of rsmA (PA0905), pilA (PA4525), tseT and tsiT (PA3907-08) | This study |
| ΔrsmA ΔpilA Δtse5_tsi5 | Deletion of rsmA (PA0905), pilA (PA4525), tse5 and tsi5 (PA2683-84) | This study |
