## Supplementary material for "Effectiveness of *Pseudomonas aeruginosa* type VI secretion system relies on toxin potency and type IV pili-dependent interaction": S2 Table

**S2 Table List of plasmids used in this study**

| **Plasmid constructs** | **Features** | **Source** |
| --- | --- | --- |
| pCR-BluntII-TOPO | Blunt sub-cloning vector, KmR | Invitrogen |
| pKNG101 | Suicide vector, Sm^R^ | Kaniga et.al. 1991 |
| pKNG101_ΔrsmN | Suicide vector to delete *rsmN* (*PA5183.1*) from *P. aeruginosa*, Sm^R^ | This study |
| pKNG101_ΔtssB1 | Suicide vector to delete *tssB1* (*PA0083*) from *P. aeruginosa*, Sm^R^ | Laboratory collection |
| pKNG101_ΔtssB2 | Suicide vector to delete *tssB2* (*PA1657*) from *P. aeruginosa*, Sm^R^ | Laboratory collection |
| pKNG101_ΔtssB3 | Suicide vector to delete *tssB3* (*PA2365*) from *P. aeruginosa*, Sm^R^ | Laboratory collection |
| pKNG101_Δtse1tsi1 | Suicide vector to delete tse1 and tsi1 (PA1844-45) from *P. aeruginosa*, SmR | Laboratory collection |
| pKNG101_Δtse2tsi2 | Suicide vector to delete tse2 and tsi2 (PA2702-03) from *P. aeruginosa*, SmR | Laboratory collection |
| pKNG101_Δtse3tsi3 | Suicide vector to delete tse3 and tsi3 (PA3484-85) from *P. aeruginosa*, SmR | Laboratory collection |
| pKNG101_Δtse4tsi4 | Suicide vector to delete tse4 and tsi4 (PA2774-75) from *P. aeruginosa*, SmR | Laboratory collection |
| pKNG101_Δtse5tsi5 | Suicide vector to delete tse5 and tsi5 (PA2683-84) from *P. aeruginosa*, SmR | This study |
| pKNG101_Δtse6tsi6 | Suicide vector to delete tse6 and tsi6 (PA1844-45) from *P. aeruginosa*, SmR | This study |
| pKNG101_Δtse7tsi7 | Suicide vector to delete tse7 and tsi7 (PA0099-0100) from *P. aeruginosa*, SmR | This study |
| pKNG101_Δtse8tsi8 | Suicide vector to delete tse8 and tsi8 (PA1844-45) from *P. aeruginosa*, SmR | This study |
| pKNG101_Δtle1tli1ab | Suicide vector to delete tle1, tli1a and tli1b (PA3290-92) from *P. aeruginosa*, SmR | Laboratory collection |
| pKNG101_Δtle3tli3 | Suicide vector to delete tle3 and tli3 (PA0259-60) from *P. aeruginosa*, SmR | Laboratory collection |
| pKNG101_Δtle4tli4 | Suicide vector to delete tle4 and tli4 (PA1509-10) from *P. aeruginosa*, SmR | Laboratory collection |
| pKNG101_ΔpldAtli5a | Suicide vector to delete pldA and tli5a (PA3487-88) from *P. aeruginosa*, SmR | Laboratory collection |
| pKNG101_ΔpldBtli5b1-3 | Suicide vector to delete pldB, tli5b1, tli5b2 and tli5B3 (PA5086-89) from *P. aeruginosa*, SmR | Laboratory collection |
| pKNG101_ΔtseTtsiT | Suicide vector to delete tseT and tsiT (PA3907-08) from *P. aeruginosa*, SmR | This study |
| pKNG101_ΔtseVtsiV | Suicide vector to delete tseV and tsiV (PA0821-22) from *P. aeruginosa*, SmR | This study |
| pKNG101_ΔvgrG2b_vgrG2bi | Suicide vector to delete vgrG2b and vgrG2bi (PA0261-62) from *P. aeruginosa*, SmR | Laboratory collection |
| pKNG101_ΔampDh3_ampDh3i | Suicide vector to delete ampDh3 and ampDh3i (PA0807-08) from *P. aeruginosa*, SmR | This study |
| pKNG101_Δazu | Suicide vector to delete azu (PA4922) from *P. aeruginosa*, SmR | This study |
| pKNG101_ΔpilA | Suicide vector to delete pilA (PA4525) from *P. aeruginosa*, SmR | This study |
| pKNG101_tssB1:mScarlet-I | Suicide vector to deliver TssB1-mScarlet-I fusion into *P. aeruginosa*, Sm^R^ | Laboratory collection |
| pKNG101_tssB2:sfGFP | Suicide vector to deliver TssB2-sfGFP fusion into *P. aeruginosa*, Sm^R^ | Laboratory collection |
| pUCP22 | Plasmid containing promoterless GFPmut3b, GmR | Tim Tolker-Nielsen, University Copenhagen |
| pUCP22_tssA1_transcrip. | Plasmid containing GFPmut3b under tssA1g transcirptional control, GmR | This study |
| pUCP22_tssA1_transl. | Plasmid containing GFPmut3b under tssA1 translational control, GmR | This study |
| pUCP22_tssA2_transcrip. | Plasmid containing GFPmut3b under tssA2 transcirptional control, GmR | This study |
| pUCP22_tssA2_transl. | Plasmid containing GFPmut3b under tssA2 translational control, GmR | This study |
| pUCP22_tssB3_transcrip. | Plasmid containing GFPmut3b under tssB3 transcirptional control, GmR | This study |
| pUCP22_tssB3_transl. | Plasmid containing GFPmut3b under tssB3 translational control, GmR | This study |
| miniCTX-pX2-mCherry | mini-CTX suicide vector cartying mCherry under pX2 promoter, TcR | Knut Drescher Lab, Biozentrum University of Basel |
| miniCTX-pX2-sfGFP | mini-CTX suicide vector cartying sfGFPunder pX2 promoter, TcR | Knut Drescher Lab, Biozentrum University of Basel |
