## Supplementary material for "Effectiveness of *Pseudomonas aeruginosa* type VI secretion system relies on toxin potency and type IV pili-dependent interaction": S3 Table

**S3 Table List of primers used in this study**

| **Name** | **Sequence** | **Description** |
| --- | --- | --- |
| **Vector primers** | | |
| UpKn | CTCATCAGTGAAATCCAGGG | Primer flanking MCS of pKNG101 |
| RpKn | CATATCACAACGTGCGTGGA | Primer flanking MCS of pKNG101 |
| M13 Fw | TGTAAAACGACGGCCAGT | M13 forward sequencing primer |
| M13 Rw | CAGGAAACAGCTATGACC | M13 reverse sequencing primer |
| OAL1604 | CGCGCGTAATACGACTCACT | Primer flanking MCS of mini-CTX plasmids |
| OAL1605 | CCGTCCTTGCTGAATTAGCTT | Primer flanking MCS of mini-CTX plasmids |
| **Construction of mutator plasmids and screening of gene deletions** | | |
| OAL3239 | ATGATGGGCCCGCTCCAGGTTGAGCTGATTGAGGC | Constructing mutator for rsmN deletion P1 (SmaI) |
| OAL3240 | CAACTCGTCGAAACCCATGTTCCGCGT | Constructing mutator for rsmN deletion P2 |
| OAL3241 | ATGGGTTTCGACGAGTTGAAGACGGCACCG | Constructing mutator for rsmN deletion P3 |
| OAL3242 | ATCATGGATCCTAATCGCGTTCGGCCTGCTG | Constructing mutator for rsmN deletion P1 (BamHI) |
| OAL6642 | GCTTGCCGAAGCTGATGTGT | Screening rsmN deletion |
| OAL6643 | CTCGACGCTGGAGCAATGTT | Screening rsmN deletion |
| OAL2661 | CGACCCCACCTTCCGTATCAAC | Screening tssB1 deletion / fluorescent fusion |
| OAL2662 | CGATGTAGCGGGAGTCCTCG | Screening tssB1 deletion / fluorescent fusion |
| OAL1594 | CAGGCGATGCGGGAAGTCGAAA | Screening tssB2 deletion / fluorescent fusion |
| OAL1595 | TCTGCCACTTGGCGAACTGC | Screening tssB2 deletion / fluorescent fusion |
| OAL3149 | GAAGGACTCCGACTCGATGAAC | Screening tssB3 deletion |
| OAL3150 | GACGTTCGACAGCTTCTCCA | Screening tssB3 deletion |
| OAL1295 | CACTTGGCGGTAAAGATCG | Screening tse1tsi1 deletion |
| OAL1296 | GATGGCCTGGATCACGTC | Screening tse1tsi1 deletion |
| OAL1283 | GCTGCTGCTCGGCCTGTTC | Screening tse2tsi2 deletion |
| OAL1284 | AGGCGCCTGAACTGTTCGT | Screening tse2tsi2 deletion |
| OAL1345 | CCATTACGCCGAACTCACC | Screening tse3tsi3 deletion |
| OAL4465 | CGGGGTGAAGGCGAGGAAGG | Screening tse3tsi3 deletion |
| OAL4649 | CCGACGCGGCGTAATAGG | Screening tse4tsi4 deletion |
| OAL4650 | CTGGATGCCCGGCCAGGC | Screening tse4tsi4 deletion |
| OAL6254 | ATTATAGTCGACCACGGGTCGCCTCGATCTTCACC | Constructing mutator for tse5tsi5 deletion P1 (SalI) |
| OAL6255 | ATGAGCGGCGACGAGTGAGGCCGACCC | Constructing mutator for tse5tsi5 deletion P2 |
| OAL6256 | TCACTCGTCGCCGCTCATCTATCCATCCTTCCTTG | Constructing mutator for tse5tsi5 deletion P3 |
| OAL6257 | ATTATAGGGCCCGCAACGAGACGGTCAGGATCGGC | Constructing mutator for tse5tsi5 deletion P4 (ApaI) |
| OAL6258 | CAGCACGTCGGGGTGATC | Screening tse5tsi5 deletion |
| OAL6259 | ACGAGACGGTGCATGTGAAG | Screening tse5tsi5 deletion |
| OAL6260 | ATTATAGGATCCCAACAAGAGCATCGGCCACGAC | Constructing mutator for tse6tsi6 deletion P1 (BamHI) |
| OAL6261 | ATGGATGCGCTGCCCTGAGCGCAACACC | Constructing mutator for tse6tsi6 deletion P2 |
| OAL6262 | TCAGGGCAGCGCATCCATGCGTCGGTAC | Constructing mutator for tse6tsi6 deletion P3 |
| OAL6263 | ATTATAGGGCCCCAGGAAAGCCTTGATTCGCGAGC | Constructing mutator for tse6tsi6 deletion P4 (ApaI) |
| OAL3352 | AGTTATACATCCACGCCGAGC | Screening tse6tsi6 deletion |
| OAL3353 | GAAAGGGGAGATGCGTGACA | Screening tse6tsi6 deletion |
| OAL6264 | ATTATAGGATCCCAACGGCTTCATTCCCGGCG | Constructing mutator for tse7tsi7 deletion P1 (BamHI) |
| OAL6265 | TGCTGGCCGTTGGCCATCAGGCAGCC | Constructing mutator for tse7tsi7 deletion P2 |
| OAL6266 | ATGGCCAACGGCCAGCAATGGACGCG | Constructing mutator for tse7tsi7 deletion P3 |
| OAL6267 | ATTATAGGGCCCGCCAAGGGCGGACAGCAG | Constructing mutator for tse7tsi7 deletion P4 (ApaI) |
| OAL3307 | TGGAAGGCGAGCTGGGAC | Screening tse7tsi7 deletion |
| OAL3308 | TGCCAGGCGAGCAGCA | Screening tse7tsi7 deletion |
| OAL6268 | ATTATAGGATCCCGAGAGCAATCCGCGCAGC | Constructing mutator for tse8tsi8 deletion P1 (BamHI) |
| OAL6269 | TCAGTCGCGCTCGATCATGCTGTCACCGCC | Constructing mutator for tse8tsi8 deletion P2 |
| OAL6270 | ATGATCGAGCGCGACTGAGCGCTTGCC | Constructing mutator for tse8tsi8 deletion P3 |
| OAL6271 | ATTATAGGGCCCCGCATCGGCTACATGGTCCTG | Constructing mutator for tse8tsi8 deletion P4 (ApaI) |
| OAL6272 | CTGCACAGGTTGACGATGC | Screening tse8tsi8 deletion |
| OAL6273 | CGACAGCGAGTTCCACTACG | Screening tse8tsi8 deletion |
| OAL4321 | GCGCTGTTCAAATTGCTGGAG | Screening tle1tli1ab deletion |
| OAL4322 | AGAACAGGCCAGGCATTCTAG | Screening tle1tli1ab deletion |
| OAL3293 | CCGGGAAAGACGTTGAAGGA | Screening tle3tli3 deletion |
| OAL3294 | GTAGGTTCGGATGGCGGTAG | Screening tle3tli3 deletion |
| OAL2586 | GTTTTCAGCGACCCCTACCTC | Screening tle4tli4 deletion |
| OAL2053 | CCGCAGCAAACCCTCCAG | Screening tle4tli4 deletion |
| OAL3231 | CCGGGCAGAAGATGGTGATC | Screening pldAtli5a deletion |
| OAL3232 | AGGACGATGCAATTGGTGGT | Screening pldAtli5a deletion |
| OAL3237 | GAACTGGCCACCTTGCATTC | Screening pldBtli5b1-3 deletion |
| OAL3238 | GGGCGACGAGATCCATTTCA | Screening pldBtli5b1-3 deletion |
| OAL6274 | ATTATAGTCGACGCGCTGGAACAGGTCATGACC | Constructing mutator for tseTtsiT deletion P1 (SalI) |
| OAL6275 | TCAGCCGCGGCCGCTCATGCCGGTCTC | Constructing mutator for tseTtsiT deletion P2 |
| OAL6276 | ATGAGCGGCCGCGGCTGACGACGTACG | Constructing mutator for tseTtsiT deletion P3 |
| OAL6277 | ATTATAGGGCCCGGCCTGTAGGGCGAATAACCG | Constructing mutator for tseTtsiT deletion P4 (ApaI) |
| OAL6278 | CATGGCCTCCTGGCTGTG | Screening tseTtsiT deletion |
| OAL6279 | GCCGGGTGGCGATGAAG | Screening tseTtsiT deletion |
| OAL6548 | ATTATAGGATCCCGAGTCTCGCACCGAGG | Constructing mutator for tseVtsiV deletion P1 (BamHI) |
| OAL6549 | ATGACCAAGCGCCTGTAGGGGCTTACG | Constructing mutator for tseVtsiV deletion P2 |
| OAL6550 | CTACAGGCGCTTGGTCATGTCGTAGCTGACC | Constructing mutator for tseVtsiV deletion P3 |
| OAL6551 | ATTATAACTAGTCCTTAACCGCAACAGCGTG | Constructing mutator for tseVtsiV deletion P4 (SpeI) |
| OAL6552 | TCACCTTTGATAGCATCCTGGC | Screening tseVtsiV deletion |
| OAL6553 | GTGACGCTGGGTAGTATCGG | Screening tseVtsiV deletion |
| OAL3320 | TGAACTAGTGTAACGCTTGCGGATGATCTTG | Screening vgrG2b vgrG2bi deletion |
| OAL3321 | CCCGACGACATTGATGGTGT | Screening vgrG2b vgrG2bi deletion |
| OAL6824 | ATTATAGGATCCAGTAGCCGCTCTGTCGAGGGT | Constructing mutator for ampDh3ampDh3i deletion P1 |
| OAL6825 | TCAGCCATCGAGACCGCGATAGCTGTTGTAGTCG  ATGGTCAGCATGG | Constructing mutator for ampDh3ampDh3i deletion P2 |
| OAL6826 | GACTACAACAGCTATCGCGGTCTCGATGGCTGAG  CATTG | Constructing mutator for ampDh3ampDh3i deletion P3 |
| OAL6827 | ATTATAACTAGTAACGCGGCGATCCTGATCGTC | Constructing mutator for ampDh3ampDh3i deletion P4 |
| OAL6671 | CCGCTGGCACGACGCTAG | Screening ampDh3 ampDh3i deletion |
| OAL6672 | ACCTGGCGATCGGCATCGT | Screening ampDh3 ampDh3i deletion |
| OAL6673 | ATTATAACTAGTTGGCCTACAACAAGGACAACCAGG | Constructing mutator for PA5264 PA5265 deletion P1 (SpeI) |
| OAL6674 | AGCGGATTCGAGCACTGATGCTGAGTTATTCCGC | Constructing mutator for PA5264 PA5265 deletion P2 |
| OAL6675 | TCAGTGCTCGAATCCGCTCATGCCTCGCTC | Constructing mutator for PA5264 PA5265 deletion P3 |
| OAL6676 | ATTATAGGATCCACATCGAGCACGACCAGAAGATCC | Constructing mutator for PA5264 PA5265 deletion P4 (BamHI) |
| OAL6677 | TCATCGACCTGCCCGACC | Screening PA5264 PA5265 deletion |
| OAL6678 | AAGACCCGCAGCGTCTTCA | Screening PA5264 PA5265 deletion |
| OAL6408 | ATTATAATGCATCCTGGAACAACTGGCCGGC | Constructing mutator for azu deletion P1 (NsiI) |
| OAL6409 | ATGCTACGTCTGAAGTGATGCGCGAGCG | Constructing mutator for azu deletion P2 |
| OAL6410 | TCACTTCAGACGTAGCATGGAGCAGCCTC | Constructing mutator for azu deletion P3 |
| OAL6411 | ATTATAGGATCCCGATCTCGGCCTCCTGCAGG | Constructing mutator for azu deletion P4 (BamHI) |
| OAL6412 | AGCTGTATCCCTGCGAAGG | Screening azu deletion |
| OAL6413 | GGCAAGGGCCAGAAGATCG | Screening azu deletion |
| OAL6508 | ATTATAGGATCCCGCATAGCACCCGGCAAG | Constructing mutator for pilA deletion P1 (BamHI) |
| OAL6509 | ATGAAAGCTGATAACTAAGGTGATCGAAGGTG | Constructing mutator for pilA deletion P2 |
| OAL6510 | CTTAGTTATCAGCTTTCATGAATCTCTCCGT | Constructing mutator for pilA deletion P3 |
| OAL6511 | ATTATAGGTACCGTGTTGGCGGACCAGCT | Constructing mutator for pilA deletion P4 (KpnI) |
| OAL6512 | GCCACAACCATCGCATCGG | Screening pilA deletion |
| OAL6513 | CGACCAGAATCGCTTCGGTC | Screening pilA deletion |
| **Construction of reporter plasmids** | | |
| OAL5919 | TAATAAGGTACCGGTCAGCTGTCCCTGGTCCAGT | Constructing transcriptional / trasnational fusion for tssA1 promoter (KpnI) |
| OAL6365 | ATTATAGGATCCGGCAGCCAGCAAAACGGGTAC | Constructing transcriptional fusion for tssA1 promoter (BamHI) |
| OAL5921 | TAATAAGGTACCGCGCAGTAGTGTTCAGGCCAGT | Constructing transcriptional / trasnational fusion for tssA2 promoter (KpnI) |
| OAL6366 | ATTATAGGATCCATGCGAGGAGAGCTTGCTCGAAT | Constructing transcriptional fusion for tssA2 promoter (BamHI) |
| OAL5923 | TAATAAGGTACCCTCCATACCGCGAACTGCTGCC | Constructing transcriptional / trasnational fusion for tssB3 promoter (KpnI) |
| OAL6367 | ATTATAGGATCCGTCCAGCTTGTGCTGCGTACTC | Constructing transcriptional fusion for tssB3 promoter (BamHI) |
| OAL6530 | TTTACGCATGGCAGCCAGCAAAACGG | Constructing translational fusion for tssA1 promoter P2 |
| OAL6531 | CTGGCTGCCATGCGTAAAGGAGAAGAACTTTTCAC | Constructing translational fusion for tssA1 promoter P3 |
| OAL6532 | GAGTCAGCTCAGCTAATAAGCTTATTTGTAT | Constructing translational fusion for tssA1 promoter P4 |
| OAL6533 | ATTATAGCATGCAATGCGAGGAGAGCTTGCTC | Constructing translational fusion for tssA2 promoter (SphI) |
| OAL6534 | ATTATAGCATGCGGTCCAGCTTGTGCTGCG | Constructing translational fusion for tssB3 promoter (SphI) |
| OAL5919 | TAATAAGGTACCGGTCAGCTGTCCCTGGTCCAGT | Constructing transcriptional / trasnational fusion for tssA1 promoter (KpnI) |
