## Supplementary material for "Effectiveness of *Pseudomonas aeruginosa* type VI secretion system relies on toxin potency and type IV pili-dependent interaction": S4 Table

**S4 Table Simulation parameters**

| **Parameter** | **Value(s)** | **Source** |
| --- | --- | --- |
| **Simulation initiation** | | |
| Inoculum radius* | 110μm |  |
| Inoculum density (OD)** | 0.1, 8.0, 4.0, 2.0, 1.0, 0.5, 0.25, 0.125 |  |
| Attacker to prey ratio in inoculum | 0.5, 0.45, 0.4, 0.35, 0.3, 0.25, 0.2, 0.15, 0.1, 0.05 |  |
| **Biophysics** | | |
| Time step | 0.025h | [1] |
| Cell growth drag | 10 | [1] |
| Max contacts | 24 | [1] |
| Number of sub-steps | 8 | [1] |
| Division orientation noise | 0.10% | [1] |
| **Cells parameters** | | |
| Cell radius | 0.5μm | Estimated from single cell microscopy images |
| Target division length | 3μm | Estimated from single cell microscopy images |
| Cell division noise | 0.5 | [2] |
| Maximum growth rate | 1 h^-1^ | [2] |
| **T6SS parameters** | | |
| T6SS firing rate | 1, 5, 25, 50, 100, 200 | [3] |
| Toxin lethal dose | 1, 5, 10, 20, 50 | [3] |
| Toxin lysis delay (min) | 3, 60, 120, >simulation time | [3] |
| Fraction of attacker population actively firing | 1, 0.5, 0.25, 0.125, 0.0625 | This study |
| Upfront T6SS cost | 0 | [4] |
| Cost per T6SS firing event | 0.00001 | [4] |
| Needle length | 0.5μm | [4] |
| Minimum needle penetration | 10nm | [3] |

* 1μL spot drying on agar will result in inoculum area of 7.55 * 10^6^ μm^2^ (Estimated from whole colony microscopy images)

** 2.04*10^5^ colony forming units per 1μL of OD600=1.0 culture as per Kim et.al. 2012
