## Supporting figure legendfs for "Effectiveness of *Pseudomonas aeruginosa* type VI secretion system relies on toxin potency and type IV pili-dependent interaction": Supporting information.docx

**S1** **Fig. Gac-Rsm mutant strains show graduated increase in H1-T6SS and H2-T6SS promoter translational activity.** Analysis of H1-T6SS (**A**) and H2-T6SS (**B**) promoter translational activity in WT, *rsmN*, *rsmA*, *retS*, and *rsmArsmN* strains. Measurements performed in static biofilm using plasmid based GFPmut3b reporter fusions. All measurements performed at 37^o^C, in static culture, each display item shows a mean +SD of 4 technical replicates that is representative of n = 3 biologically independent repeats.

**S2 Fig.** **Gac-Rsm mutant strains show graduated increase in H3-T6SS promoter activity and expression.** Analysis of H3-T6SS promoter transcriptional (**A**) and translational (**B**) activity in WT, *rsmN*, *rsmA*, *retS*, and *rsmArsmN* strains. Measurements performed in static biofilm using plasmid based GFPmut3b reporter fusions. All measurements performed at 37^o^C, in static culture, each display item shows a mean +SD of 4 technical replicates that is representative of n = 3 biologically independent repeats. (**C**) Western blot analysis shows gradual elevation in of Hcp3 expression in WT, *rsmN*, *rsmA*, *retS*, and *rsmArsmN* strains. A representative blot of 3 independent biological repeats shown here, for H3-T6SS activity assessment bacteria were cultured for 24h at 30^o^C. RNA polymerase (RpoB) used as loading control.

**S3 Fig. Decrease in growth temperature results in elevated H2-T6SS and H3-T6SS promoter activity, while H1-T6SS promoter activity is decreased in Gac-Rsm regulatory cascade mutant strains.** Analysis of T6SS promoter transcriptional (left) and translational (right) activity in WT, *rsmN*, *rsmA*, *retS*, and *rsmArsmN* strains. Upper row shows H1-T6SS (*tssA1*), middle H2-T6SS (*tssA2*), and lower H3-T6SS (*tssB3*), promoter activity over growth time as measured by plasmid based GFPmut3b reporter fusions. All measurements performed at 25^o^C, in static culture, each display item shows a mean +SD of 4 technical replicates that is representative of n = 3 biologically independent repeats.

**S4 Fig. Changes in distribution of individual sub-populations in macrocolonies of Gac-Rsm mutants depending on inoculum density.** Single channel images corresponding to composite in Fig. 2A – mCherry in red (**A**) and sfGFP in green (**B**). Mixed bacterial colonies of WT, *rsmA*, *retS*, and *rsmArsmN* strains with altered inoculum densities. Isogenic bacterial strains tagged with mCherry (red) and sfGFP (green) fluorophores were mixed at 1 to 1 ratio and after adjusting inoculum density (OD_600_ 1.0; 0.1; 0.01; 0.001) spotted on LB agar, images of whole microcolonies taken after 48h incubation at 37^o^C show 2 morphologically distinct regions - highly mixed inner region corresponding to inoculum zone and outer region where spatial segregation of the sub-populations is apparent.

**S5 Fig. Increase in fluorescence signal variance and decreased intermixing are indicative of sub-population segregation in outer colony regions.** Relative mean signal intensity + SD of individual fluorescence channels (**A**) and relative intermixing (**B**) are calculated for circular sections taken at increasing distance from colony central point. Calculations correspond to fluorescence images shown in Fig. 2.

**S6 Fig. Loss of the ability to fire T6SS does not result in loss of competitive fitness in isogenic strains.** Fluorescent images of whole bacterial colonies of bacteria mixes made up of T6SS+ (*retS*) and T6SS– (*retStssB1tssB2tssB3*) strains at 1:1 initial ratio and initial OD_600_ 1.0, showing individual channel and overlay images of 3 biological repeats.

**S7 Fig. Further elevation in H1-T6SS activity enables further toxin sensitised prey elimination.** Representative image of 48h old mixed colonies of toxin sensitised bacteria in red in competition with T6SS+ or T6SS- (*tssB1tssB2tssB3*) strains of the same regulatory background in green. Upper lane contains a control mix of bacteria with full toxin-immunity gene sets, each of the following lanes contains strain sensitised to one of the H1-T6SS toxins from Tse1 to Tse8. Images sets of competitions of *retS* (**A**) and *rsmArsmN* (**B**) background strains shown with each of the sets containing both single fluorescence channel and overlay images showing distribution of sensitised prey in a mix with T6SS^+^ and subsequently T6SS- (*tssB1tssB2tssB3*) parental strain. Strains contain constitutively expressed fluorescent proteins, prey labelled with mCherry (shown in red) and attacker with sfGFP (shown in green). All bacteria mixed at 1:1 ratio, inoculum OD_600_ 1.0, grown for 48h at 37^o^C on LB with 2% (w/v) agar.

**S8 Fig. H2-T6SS toxin mediated killing is observed when bacteria are grown at reduced temperature.** Sensitivity to H2-T6SS toxins can be observed when growing colonies at 25^o^C, but not 37^o^C. Representative images of 48h old mixed colonies of toxin sensitised *rsmA* bacteria in red in competition with T6SS+ (*rsmA*) or T6SS- (*rsmAtssB1tssB2tssB3*) parental strain in green. Upper lane contains a control mix of bacteria with full toxin-immunity gene sets, each of the following lanes contains strain sensitised to one of the H2-T6SS toxins in the following order: Tle1, Tle3, Tle4, PldA, PldB, TseT, TseV, VrgG2b, AmpDh3, PA5265, and common good effector Azu. Image sets of competitions show both single fluorescence channel and overlay images depicting distribution of sensitised prey in a mix with T6SS^+^ and subsequently T6SS- (*tssB1tssB2tssB3*) parental strains at 2 different growth temperatures – 37^o^C and 25^o^C. Strains contain constitutively expressed fluorescent proteins, prey labelled with mCherry (shown in red) and attacker with sfGFP (shown in green). All bacteria mixed at 1:1 ratio, inoculum OD_600_ 1.0, grown for 48h, at 37^o^C bacteria incubated on LB with 2% (w/v) agar, and at 25^o^C bacteria incubated on LB with 1.2% (w/v) agar.

**S9 Fig. Increased H2-T6SS toxin mediated prey growth restriction can be observed in *retS* strains.** Representative images of 48h old mixed colonies of toxin sensitised *retS* bacteria in red in competition with T6SS+ (*retS*) or T6SS- (*retStssB1tssB2tssB3*) parental strain in green. Upper lane contains a control mix of bacteria with full toxin-immunity gene sets, each of the following lanes contains strain sensitised to one of the H2-T6SS toxins in the following order: Tle1, PldA, PldB, TseT and AmpDh3. Image sets of competitions show both single fluorescence channel and overlay images depicting distribution of sensitised prey in a mix with T6SS+ and subsequently T6SS- (*tssB1tssB2tssB3*) parental strains. Strains contain constitutively expressed fluorescent proteins, prey labelled with mCherry (shown in red) and attacker with sfGFP (shown in green). All bacteria mixed at 1:1 ratio, inoculum OD_600_ 1.0, grown for 48h at 25^o^C on LB with 1.2% (w/v) agar.

**S10 Fig. Inoculum composition determines prey/attacker contact assortment within macrocolony. (A)** Representative simulation outputs showing changes species distribution resulting from variation in initial density and mixing ratio of the populations. **(B)** Changes in localised agent intermixing as interspecies contact assortment resulting from variation in initial density and mixing ratio of cells in simulation setup. (No T6SS interactions, mean of n=5).

**S11 Fig. Time resolved changes in interspecies contacts are determined by both inoculum density and initial ratio of strains in the mix. (A, D)** Total number of cells over simulation time-course. **(B, E)** Total number of contact cells over simulation time-course. **(C, F)** Interspecies contact assortment over 30simulation time-course. **A, B, and C** effect of variation in initial density. **(D, E, F)** effect of variation in species mixing ratio. (No T6SS interactions, mean +SD of n=5).

**S12 Fig. Set of simulation outputs (A) corresponding to Fig. 5C with non-lytic toxin-based interactions. Set of corresponding green-fluorescent channel images (B) for the Fig. 5D and a set of images of preys from Fig. 5D in a mix with an T6SS- attacker strain (C). (A)** Representative simulation outputs showing how decrease in initial bacterial density promotes prey (red) survival in a mix an attacker population (green) with differing mean T6SS firing rate within a context of non-lytic toxins. (Toxin lethal dose = 5). **(B)** Single channel images corresponding to composite in Fig. 5D showing distribution of attacker population in green. **(C)** Single channel and composite images of the prey set from Fig. 5D. In a mix with T6SS- attacker (*tssB1tssB2tssB3*) of corresponding regulatory background. Mixed bacterial colonies of WT, *rsmA*, *retS*, and *rsmArsmN* strains with altered inoculum densities. Isogenic bacterial strains tagged with mCherry (red) and sfGFP (green) fluorophores were mixed at 1 to 1 ratio and after adjusting inoculum density (OD 1.0; 0.1; 0.01; 0.001) spotted on LB agar, images of whole microcolonies taken after 48h incubation at 37^o^C on LB with (2% w/v) agar.

**S13 Fig. Set of corresponding green-fluorescent channel images (A) for the Fig. 5E and a set of images of preys from Fig. 5E in a mix with an T6SS- attacker strain (B). (A)** Fluorescent single (sfGFP) channel images of mixed colonies from Fig. 5F showing distribution of T6SS+ attacker strains. **(B)** Tse5 sensitive prey spatial distribution in presence of T6SS**-** *(tssB1tssB2tssB3)* parental competitor strain. Individual and overlaid channel images shown. Each of the columns correspond to WT, *rsmA*, *retS*, and *rsmArsmN* regulatory background strains in the given order. Each of the rows contains a set of representative images of colonies set up with a different prey to attacker ratio in the inoculum. Prey to attacker ratios from the top row are as follows: 1:1, 2:1, 3:1, 5:1, 10:1. (Strains contain constitutively expressed fluorescent proteins, prey labelled with mCherry (shown in red) and attacker with sfGFP (shown in green). All bacteria mixed at ratios specified, inoculum OD_600_ 0.01, grown for 48h at 37^o^C on LB with 2% (w/v) agar.

**S14 Fig. Loss of T4P interferes with H1-T6SS mediated killing in the same manner as H2-T6SS.** Tse5 mediated competition was used to assess impact on H1-T6SS mediated killing in a mix of T4P+ and T4P- bacteria. The panel contains individual red channel images showing toxin sensitive prey distribution in presence of T6SS+ (left) or T6SS- (right) attacker strain in the upper two rows. Corresponding overlay images showing distribution of both toxin sensitive prey (in red - mCherry) and T6SS attacker population in green (sfGFP) shown in the lower two rows. T4P+ prey (*rsmA tse5tsi5*) strains in the upper row, with T4P- prey (*rsmA pilA tse5tsi5*) in the lower row. With T4P+ attacker in the first column and T4P- attacker strains in the second.

All bacteria mixed at 1 to 1 ratio, with inoculum density OD_600_ 1.0, grown for 48h at 37^o^C on LB with 2% (w/v) agar. 1 of 1 biological repeat shown for all images containing *pilA* mutant strains.

**Video 1. H1-T6SS dynamics is *rsmA* deletion strain.** Gradual sheath assembly and firing visualised using *tssB1*::mScarlet-I chromosomal fusion at native locus of *rsmA* mutant bacteria growing on agar surface. The field of view of approx. 15.6 x 13 µm, showing 30 timelapse frames captured every 4s.

**Video 2. H1-T6SS dynamics is *retS* deletion strain.** Gradual sheath assembly and firing visualised using *tssB1*::mScarlet-I chromosomal fusion at native locus of *retS* mutant bacteria growing on agar surface. The field of view of approx. 15.6 x 13 µm, showing 30 timelapse frames captured every 4s.

**Video 3. H1-T6SS dynamics is rsmA rsmN double deletion strain.** Gradual sheath assembly and firing visualised using *tssB1*::mScarlet-I chromosomal fusion at native locus of *rsmA* and *rsmN* mutant bacteria growing on agar surface. The field of view of approx. 15.6 x 13 µm, showing 30 timelapse frames captured every 4s.

**S1 Table List of strains used in this study**

**S2 Table List of plasmids used in this study**

**S3 Table List of primers used in this study**

**S4 Table Simulation parameters**
